# Maximising average mechanical power output during stretch-shortening cycles of rat medial gastrocnemius muscle

**DOI:** 10.1101/2025.09.25.678481

**Authors:** Edwin D.H.M. Reuvers, Huub Maas, Wendy Noort, Maarten F. Bobbert, Dinant A. Kistemaker

## Abstract

The average mechanical power output (AMPO) during a stretch-shortening cycle produced by a muscle depends on muscle length and stimulation over time. While the effects of cycle frequency and muscle length excursion on AMPO are well-known, several questions remain about the effects of muscle length and stimulation over time on the maximal attainable AMPO. For example, which precise muscle length and stimulation over time yield maximal AMPO? *In situ* experiments are inherently limited to a finite set of muscle length and stimulation over time. To overcome this limitation, we combined *in situ* experiments on rat m. gastrocnemius medialis with Hill-type muscle modelling. We first performed dedicated trials to estimate the muscle-tendon-complex (MTC) properties of each rat. Subsequently, we performed various stretch-shortening cycles with substantial differences in cycle frequency, shortening-to-lengthening time ratio and MTC length excursion. Model-predicted AMPO correlated nearly perfect with experimentally measured AMPO (r2 > 0.98). This justified further exploration using the Hill-type MTC model. Using the Hill-type MTC model, we predicted that AMPO peaks at a cycle frequency of 3.5 Hz, with a shortening-to-lengthening time ratio of 6:1 and an MTC length excursion of 8 mm. Notably, cycle frequency and MTC length excursion showed a strong interaction: increasing one necessitated a decrease in the other to maximise AMPO. By contrast, the optimal shortening-to-lengthening time ratio remained remarkably constant across all tested combinations of cycle frequency and MTC length excursion. This shows that muscles should spend substantially more time shortening than lengthening to maximise AMPO.

## 1 Introduction

In cyclic movements such as walking, running, and cycling, muscles periodically shorten and lengthen. Muscle length changes can be influenced by the individual, who can adjust — for example — frequency and amplitude within certain limits. Additionally, equipment may substantially influence how movements are executed. For instance, in bicycles the cranks are mechanically coupled, causing the legs to move in anti-phase which means that the downstroke duration of one leg equals the upstroke duration of the other. Consequently, muscle shortening and lengthening durations are more or less equal during cycling. However, it is far from trivial that equal muscle shortening and lengthening duration are optimal for maximal performance. Which ratio of muscle shortening to lengthening duration maximises performance? Or, taking it a step further, what is the optimal movement of the leg for maximal performance? To answer such questions, it is crucial to understand how muscle length over time affects the mechanical power output of muscles.

To study the mechanical output during a stretch-shortening cycle (SSC), researchers use an experimental setup in which a motor imposes cyclical length changes on either isolated muscle fibres or the entire muscle–tendon complex (MTC). While the approach is the same at both levels, this paper focuses on SSCs at the MTC level. During each SSC, the MTC first undergoes lengthening and then shortens back to its initial length. Muscle stimulation is applied during part of the SSC, while MTC force is measured. When MTC force is plotted against MTC length (change), the area enclosed by the resulting loop represents the net mechanical work produced during a full cycle. This net mechanical work is the sum of the positive mechanical work during MTC shortening (Figure 1A) and the negative mechanical work during MTC lengthening (Figure 1B). This method has been coined the ‘work loop technique’ (Josephson, 1985). The average mechanical power output (AMPO) for each cycle can then be calculated by dividing the net mechanical work per cycle by the duration of the cycle. As such, the work loop technique provides a powerful tool to study the effect of MTC length and stimulation over time on the maximal attainable AMPO of muscles.

**Figure 1:**
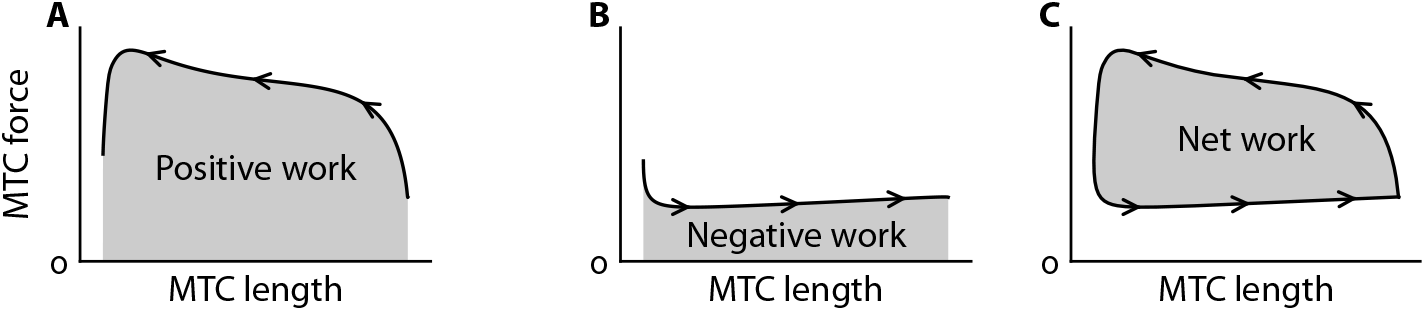
An example of a work loop. In a work loop, MTC force is plotted against MTC length (change). The area enclosed by the work loop represents the net mechanical work produced during a full cycle (C), which is the sum of the positive mechanical work during MTC shortening (A) and the negative mechanical work during MTC lengthening (B). The arrows indicate the direction of the work loop over time.

The work loop technique has been extensively employed to examine how MTC (or muscle fibre) length and muscle stimulation over time affect the mechanical behaviour of muscles. In most experiments, SSCs have been studied using sinusoidal length changes (e.g. Josephson, 1985). These SSCs are typically parametrised in terms of cycle frequency and MTC/muscle fibre length excursion, with length changes centred around the length yielding maximal isometric force. Furthermore, muscle stimulation is typically parametrised in terms of stimulation onset time and stimulation duration (i.e., ‘duty cycle’). Such parametrisation has been used in experiments to systematically investigate both the individual effects of these parameters and their interactions on AMPO. For example, results of such experiments have demonstrated a substantial interaction between cycle frequency and MTC/muscle fibre length excursion, whereby maximising AMPO typically requires larger length excursions at lower frequencies and smaller length excursions at higher frequencies (e.g., Altringham and Johnston, 1990; Caiozzo and Baldwin, 1997; James et al., 1995). Although sinusoidal SSCs have experimentally provided valuable insights, they inherently impose equal shortening and lengthening durations, leaving unexplored how unequal shortening and lengthening durations affect AMPO.

Although sinusoidal SSCs have experimentally provided valuable insights, they inherently impose equal shortening and lengthening durations. As such, the effect of unequal shortening and lengthening on AMPO remains unexplored when using sinusoidal SSCs.

Askew & Marsh (1997; 1998) performed ground-breaking experiments on the effect of unequal shortening and lengthening durations on the maximal attainable AMPO — remaining, to our knowledge, the only researchers to do so. In their studies, mouse m. soleus and m. extensor digitorum longus were connected at the distal part of the muscle belly to a servomotor for precise control of the imposed SSCs. The SSCs used by Askew & Marsh were parameterised using three variables (see Figure 2): cycle frequency, FTS (fraction of the cycle time spent shortening), and muscle fibre length excursion, with near-constant shortening and lengthening velocities. Askew & Marsh (1997; 1998) examined three distinct FTS values — 0.25, 0.50, and 0.75 — and heuristically optimised muscle fibre length excursion, muscle stimulation onset time and muscle stimulation duration to achieve the highest AMPO at different combinations of cycle frequency and FTS. Their findings indicated that AMPO was highest for FTS = 0.75, followed by FTS = 0.50, and lowest for FTS = 0.25 at each cycle frequency at the optimal muscle fibre length excursion and muscle stimulation.

**Figure 2:**
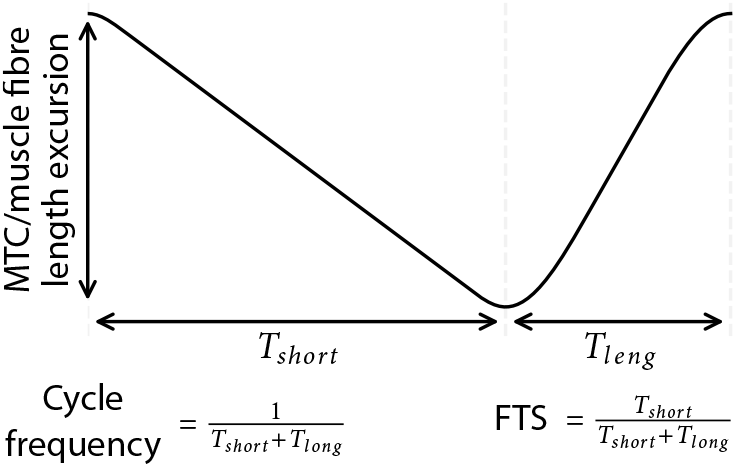
Representation of the parameterisation of stretch-shortening cycles. *T*_*short*_ and *T*_*leng*_ denote the shortening and lengthening durations, respectively, of either the muscle-tendon-complex (MTC) or the muscle fibres. FTS denotes the fraction of the cycle time spent shortening. In the example shown, the MTC/muscle fibres shorten 65% of the cycle duration (i.e., FTS = 0.65).

While Askew & Marsh’s experimental studies provided valuable insights, several questions remain regarding the effect of SSC parameters on the maximal attainable AMPO. First, Askew & Marsh (1997; 1998) optimised muscle fibre length excursion for each cycle frequency at only three distinct FTS values. Therefore, it remains unclear how muscle fibre length excursion affect the maximal attainable AMPO at different FTS values. More generally, it remains unclear how sensitive the maximal attainable AMPO is to variations in SSC parameters. For instance, does AMPO substantially decrease with suboptimal SSC parameters or is there a wide range of SSC parameter combinations that yield near-maximal AMPO? Second, because Askew & Marsh (1997; 1998) only used three distinct FTS values, it remains unclear how cycle frequency and muscle fibre length excursion affect the optimum value of FTS. More generally, it remains unclear how the optimum SSC parameters depend on each other. Third, Askew & Marsh (1997; 1998) imposed near-constant muscle fibre shortening and lengthening velocities. Consequently, it remains unclear how much more AMPO the muscle could produce if the muscle fibres were allowed to shorten-and-lengthen at their optimum. For this, the optimal muscle fibre length and stimulation over time should be identified in a parameter-free manner.

While it is obvious that muscle fibres should maximise positive mechanical work and minimise negative mechanical work in order to maximise AMPO, it is far from trivial what SSC and muscle stimulation over time yields maximal AMPO. To minimise negative mechanical work, stimulation should cease well before lengthening begins (because deactivation time delays) and muscles fibres should ideally lengthen at very low velocities (as muscle fibre force increases with increasing lengthening velocity). However, such strategies are suboptimal for AMPO: the time spent for lengthening cannot be used to produce positive mechanical work, and early deactivation reduces the positive mechanical work during shortening. One might argue that muscle fibres should shorten at a velocity that maximises the instantaneous mechanical power output to maximise positive mechanical work. However, muscle fibre force also depends on muscle fibre length, with muscle fibre force decreasing at both shorter and longer lengths than optimum length. Therefore, to maximise AMPO, it may be beneficial for muscle fibres to contract at a lower velocity than the velocity which maximises instantaneous mechanical power output. Doing so limits the negative effects of operating at suboptimal muscle fibre lengths. This raises several open questions: What shortening velocity maximises AMPO? Over what length range should muscle fibres operate? And how much time should be spent on shortening (i.e., maximising positive mechanical work) versus lengthening (i.e., minimising negative mechanical work) in order to maximise AMPO?

Obviously, it is hard to experimentally find the optimal combination of cycle frequency, FTS, MTC length excursion and muscle stimulation for maximal AMPO. If one would change MTC length excursion, how should the other SSC parameters and muscle stimulation be changed to maximise AMPO? Or, taking it one step further: How should we find the MTC length and stimulation over time that maximises AMPO when the muscle length over time is entirely free to take any form? For this reason, we propose to complement *in situ* experiments with simulations and optimisations using a Hill-type MTC model to study the effect of MTC length and muscle stimulation over time on the maximal attainable AMPO. If the MTC model yields accurate predictions of the influence of SSC parameters and muscle stimulation over time on the maximal attainable AMPO, it would allow us to answer a plethora of questions regarding the maximal attainable AMPO of various SSCs. For instance, we could investigate how the optimum SSC parameters depend on each other, and how sensitive the maximal attainable AMPO is to variations in SSC parameters. Furthermore, using an accurate model, we could optimise MTC length over time for maximal AMPO and quantify how much more AMPO could be produced compared to sinusoidal SSCs or the near-constant velocity SSCs of Askew & Marsh.

In this study, we aimed to explore the influence of SSC parameters — cycle frequency, FTS and MTC length excursion — on the maximal attainable AMPO of a single isolated MTC. As discussed earlier, only a limited set of SSCs can be investigated experimentally, so this raises the question: Which SSCs should be selected? To make an informed selection of SSCs to be tested experimentally, we first used a Hill-type MTC model — with parameter values obtained from literature — to simulate a broad range of SSCs. Based on these simulation results, we selected a smaller, representative and informative set of SSCs that captured the most prominent effects, providing a focused basis for evaluating model predictions. This set of SSCs was then tested in an *in situ* experiment on rat m. gastrocnemius medialis. In this experiment, we also performed dedicated trials to estimate the MTC properties of each specimen individually. This allowed us to compare the experimental results with model-based predictions, which showed that measured and model-predicted AMPO correlated nearly perfect (r^2^ > 0.98). The strong correspondence then justified the use of the Hill-type MTC model to (finally) explore how SSC parameters — cycle frequency, FTS and MTC length excursion — affect the maximal attainable AMPO and to identify the fully unconstrained optimal MTC length over time for maximal AMPO.

## 2 Methods

We combined *in situ* experiments with simulations and optimisations using a Hill-type MTC model to investigate the influence of SSC parameters — cycle frequency, FTS, and MTC length excursion — and muscle stimulation over time on the maximal attainable AMPO of rat m. gastrocnemius medialis (GM). We first used a Hill-type MTC model with parameter values from literature to simulate a broad range of SSCs (see Section S1). Based on these simulation results, we selected a representative and informative subset for an *in situ* experiment, in which MTC length and stimulation over time were under full control (see Section 2.1.6). In this experiment, we also performed dedicated trials to estimate the MTC properties of each specimen individually (see Section 2.1.5). Model-based predictions were then evaluated against the experimental data (see Section 2.2.1). The strong correspondence (r^2^ > 0.98) justified the use of the Hill-type MTC model to explore the influence of SSC parameters on the maximal attainable AMPO across a broad SSC parameter space (see Section 2.2.2). First, we derived predictions for SSCs with constant MTC shortening and lengthening velocity. Second, we identified the optimal SSC without any constraints on MTC length over time to investigate how much more AMPO the muscle could produce if the muscle was allowed to shorten-and-lengthen optimally (see Section S3).

### 2.1 In situ experiment

#### 2.1.1 Animals and ethics

All surgical and experimental procedures conducted in this study were approved by the Committee on the Ethics of Animal Experimentation at the Vrije Universiteit (Permit Number: FBW-AVD11200202114471) and adhered to Dutch law concerning the guidelines and regulations of animal welfare and experimentation. Data was obtained data from three male Wistar rats (body mass 365, 410 and 407 gram respectively). The rats were anaesthetised by intraperitoneally injecting a urethane solution (12.5% urethane) at a dosage of 1.2 mL/100 g body mass. Additional doses of urethane (0.2 mL/g body mass) were administered throughout the experiment as needed to maintain deep anesthesia. Heart rate, body temperature and withdrawal reflexes to a pain stimulus were monitored every 10 minutes. To prevent dehydration, subcutaneous injections of 1 mL NaCl solution (0.9%) were administered every 1-2 hours. After completion of the experiment, we killed the rats by intracardially injecting pentobarbital sodium (Euthasol 20%) followed by a double-sided pneumothorax.

#### 2.1.2 Surgical procedure

The hindlimb of the rat was shaved, and the skin was cut and partially removed. Next, m. biceps femoris was removed to expose the femur. To minimise mechanical effects of surrounding muscles on GM — in particular myofascial force transmission — the following steps were taken (see Rijkelijkhuizen et al., 2005): 1) GM was exposed as much as possible from its surrounding tissue. 2) The tendons of m. plantaris, m. gastrocnemius lateralis and m. soleus were cut at their distal ends. 3) The medial and lateral part of m. gastrocnemius were carefully separated. N. peroneus, n. suralis, and the branch of the n. tibialis innervating m. gastrocnemius lateralis and m. soleus were then cut to further reduce the influence of surrounding tissue on GM. Subsequently, the tissue around the calcaneal tendon was removed such that only a small part of the calcaneus was left. Lastly, a cuff-electrode was placed on n. ischiadicus. The part of n. ischiadicus proximal to the cuff-electrode was then crushed to prevent muscle excitation via spinal reflexes.

#### 2.1.3 Experimental setup

The rats were placed on an electrical heating pad to maintain body temperature at 37° C. The laboratory temperature was kept constant at 20 °C, with a local humidity around the muscle tissue of about 80%. A servomotor (Aurora 309C, Aurora Scientific, Aurora, Canada) was used to impose MTC length changes and to measure GM force. Before the experiment, we calibrated the position sensor of the servomotor using a dial gauge (2046S, Mitutotyo Nederland BV, Veenendaal, the Netherlands) and calibrated the force sensor using various calibration masses. To relate servomotor position (changes) to MTC length, we measured MTC length at one position with a caliper at the end of the experiment. A direct current stimulator was used to stimulate n. ischiadicus with pulse width of 100 *µ*s and at a frequency of 100 Hz during all experiments, while the (supramaximal) current was chosen for every rat individually (see Section 2.1.4). Before the series of experiments, we ensured that the position, force, and stimulation signals were synchronised in time by applying a physical stimulus that produced a detectable response across all channels. All data were collected at a sample rate of 2000 Hz.

The femur was fixed with a metal clamp and the foot with a plastic plate, such that the hindlimb was rigidly secured to the setup. The distal end of the calcaneal bone was tightly attached to a steel rod with 100% polyester yarn at the leftover part of the calcaneus. The other end of the steel rod was then attached with Kevlar thread to the motor. A similar procedure was applied to m. gastrocnemius lateralis (GL) and m. plantaris (PL). The distal tendon of GL and PL (GL+PL) was then connected to a second motor to monitor their combined force throughout the experiment in order to verify the absence of mechanical interaction with GM. Care was taken to ensure that the direction of GM force reflected *in vivo* conditions. Lastly, we connected the cuff-electrode to the direct current stimulator (DS2A, Digitimer, Welwyn Garden City, United Kingdom). This setup allowed us to have full control over MTC length and stimulation while measuring GM force (see Figure 3).

**Figure 3:**
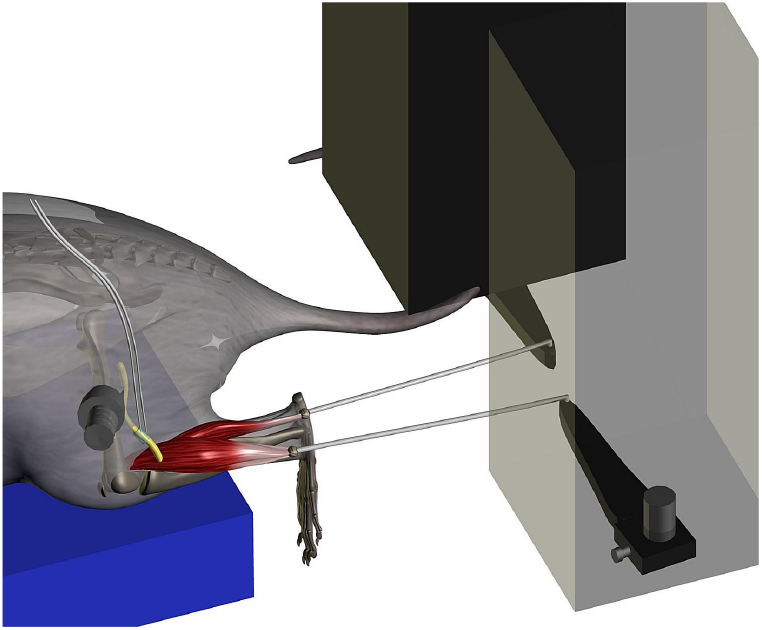
Representation of the experimental setup that provided full control of MTC length and stimulation while measuring m. gastrocnemius medialis (GM) force. GM was carefully exposed from its surrounding tissue and positioned in the setup such that GM pulled in its natural direction, while the femur and foot were securely fixated. The distal end of the calcaneal tendon was connected to a motor via a steel rod. The distal tendon of m. gastrocnemius lateralis and m. plantaris was connected to a second motor. A cuff-electrode was placed on n. ischiadicus. N. peroneus, n. suralis and the branch of n. ischiadicus innervating m. gastrocnemius lateralis and m. soleus were cut such that only GM was innervated.

#### 2.1.4 General procedures

After securing the rat in the experimental setup, we conducted a series of tetanic contractions for several purposes. First, we performed isometric contractions to ensure that GM pulled in its natural direction. These isometric contraction were also use to determine the supramaximal stimulation current for GM. The stimulation current was gradually increased until GM force plateaued. The current at which no further increase in GM force was observed was then used consistently throughout the experiment. Additionally, the isometric contractions served as a check to confirm that the surgery was successful, particularly ensuring that no GL+PL force was transmitted to the distal GL+PL tendon, and thus that no myofascial force transmission occurred (Finni et al., 2023). Subsequently, we performed two to three quick-release, step-ramp and isometric experiments along with two to three SSCs. This approach was based on results from pilot experiments, which indicated that imposing these contractions prior to the start of the experiment resulted in more stable GM force throughout the entire experiment.

Following the preparatory procedures, we commenced the main experiment. The main experiment consisted of two parts. In the first part, we performed quick-release, step-ramp and isometric experiments to estimate the MTC properties of each rat. In the second part, we performed various SSCs. To assess if GM was fatigued, two quick-release and step-ramp experiments were repeated halfway through the second part of the experiment and again at the end of the entire experiment. Additionally, isometric experiments were performed after every six series of SSCs during the second part of the experiment, using a fixed MTC length 2-4 mm below MTC optimum length. Between contractions, GM was returned to slack length, followed by a two-minute rest period. During these rest periods, GM was irrigated with saline to keep the muscle tissue moist. To reduce potentiation effects, two twitches were applied 1.5 seconds before each contraction.

#### 2.1.5 Quick-release, step-ramp and isometric experiments to estimate MTC properties

Quick-release, step-ramp, and isometric experiments were performed, with the resulting data being used to estimate MTC properties for each rat individually (see Figure 3 of Reuvers and Kistemaker, 2025). This approach served two purposes. First, it enabled us to predict AMPO for the experimentally tested SSCs for each rat individually. As a result, less rats were required to evaluate predictions against experimental results. Second, it enabled us derive predictions of the influence of a broad range of SSC parameters — cycle frequency, FTS and MTC length excursion — on the maximal attainable AMPO for each rat individually. The full experimental procedure of the quick-release, step-ramp and isometric experiments as well as the MTC property estimation has been described in Reuvers and Kistemaker (2025), where we demonstrated that these properties can be estimated accurately when accounting for muscle fibre shortening quick-releases.

#### 2.1.6 Stretch-shortening cycles to evaluate predictions

SSCs were parametrised by cycle frequency, FTS, and MTC length excursion (see Figure 2). The average MTC length was set ∼3 mm below the MTC length yielding maximal isometric force to prevent irreversible damage to the muscle fibres. MTC velocity was constant during shortening and lengthening phases, except around the transition from MTC lengthening to shortening and vice versa. During these transitioning phases, MTC length changed sinusoidally such that MTC length over time was continuously differentiable to the third order. The maximum MTC acceleration during these transitions was limited to 1500 mm/s2 to further reduce the risk of irreversible muscle fibre damage. This constraint limited the range of feasible experimental conditions: SSCs with a high cycle frequency, large MTC length excursion, and either a substantially high or low FTS value could not be tested.

Experimental conditions were selected based on preliminary simulations using a Hill-type MTC model with parameter values from literature (seem Section S1). We tested 26 conditions, comprising 13 combinations of cycle frequency and FTS (see Figure 4), each at at an MTC length excursion of 4 mm and 8 mm. Full details, including muscle stimulation onset time and duration, are provided in the Supplementary Material (Table S1 & Table S2).

**Figure 4:**
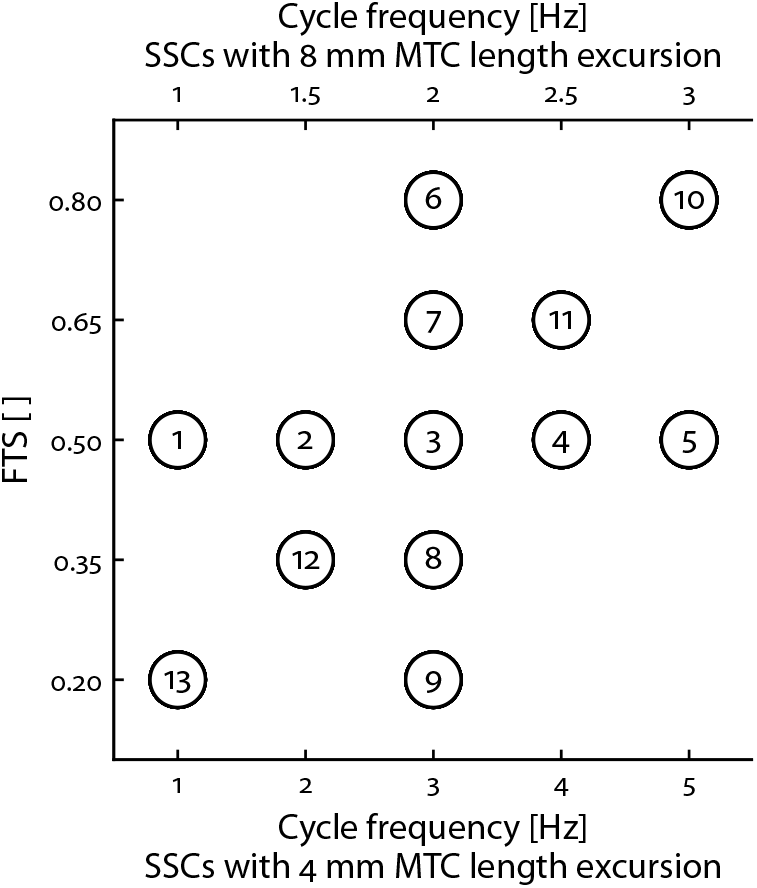
Representation of experimentally investigated SSCs. Thirteen combinations of cycle frequency and FTS were tested at two distinct MTC length excursions (4 mm and 8 mm), resulting in a total of 26 experimental SSC conditions.

Each SSC condition consisted of five consecutive cycles, with muscle stimulation applied during the middle three cycles. Muscle stimulation onset time was set at the transition from MTC lengthening to shortening. Muscle stimulation duration was based on preliminary model predictions (see Section S1), however, we chose a stimulation duration that was slightly shorter (20 ± 10 ms) than that predicted to be optimal for AMPO. This was done to prevent irreversible damage to the muscle fibres as a possible result of substantial GM force during active GM lengthening. After each SSC condition, we inspected GM force over time to heuristically adjust muscle stimulation duration to further enhance AMPO. The SSC condition was then repeated with the adjusted muscle stimulation duration after all other conditions had been performed for that specific MTC length excursion (∼40 minutes later in time). This approach aimed to evaluate to what extent we could enhance AMPO and to evaluate the reliability of measured GM force. Overall, we performed 52 series of SSCs for each rat. For each cycle, AMPO was computed as the product of cycle frequency and the integral of measured GM force over measured MTC length (i.e., the ‘work loop’ area).

Unfortunately, rat 3 passed away near the end of the experiment, which meant that six SSC conditions with an 8 mm MTC length excursion were performed only once instead of twice for this rat, such that 46 instead of 52 experimental SSCs were measured for rat 3.

### 2.2 MTC model

A Hill-type MTC model was used to represent GM. The model has been described extensively elsewhere (Reuvers and Kistemaker, 2025; Soest and Bobbert, 1993). In short, the MTC modelled consisted of a contractile element (CE) and a parallel elastic element (PEE) which were both in series with a serial elastic element (SEE). PEE and SEE were modelled as purely passive elastic structures (Zajac, 1989), with their force non-linearly dependent on their length. The behaviour of CE was more complex: CE force depended on CE length, CE velocity and active state (*q*), which was defined as the relative amount of *Ca*^2+^ bound to troponin C (Ebashi and Endo, 1968). The excitation dynamics were modelled according to Hatze (1981, pp 31-32). Here, active state depended on the normalised concentration free *Ca*^2+^ between the filaments (*γ*) and CE length as sensitivity to *Ca*^2+^ increases with CE length (e.g. Rack and Westbury, 1969; Stephenson and Williams, 1982). The concentration free *Ca*^2+^, in turn, was related to the normalised muscle stimulation (*STIM*) through a first-order differential equation. The independent inputs of the model were *STIM* and MTC length over time, while CE length and *γ* were the states.

#### 2.2.1 Evaluation of measured versus predicted AMPO

Experimental measurements were compared with predictions derived using a Hill-type MTC model. To ensure a fair comparison, MTC properties were not tuned to the experimental SSC data; instead, they were obtained from dedicated trials (see Reuvers and Kistemaker, 2025). Moreover, all model inputs were identical to those measured experimentally. Specifically, we simulated all experimental SSC conditions for each rat individually using on their estimated MTC properties, using two inputs: the experimentally measured MTC length trajectory and *STIM(t)* derived from the measured direct current stimulator signal. The latter was done because the measured direct current stimulator signal consisted of a train of pulses. *STIM* was computed as follows: *STIM* was maximal (i.e., a value of 1) between two stimulation pulses and was zero exactly halfway the inverse of the stimulation frequency (i.e., 5 ms) after the last stimulation pulse. Model-predicted AMPO was then compared with measured AMPO to evaluate predictive accuracy using the coefficient of determination (r^2^) derived from Pearson’s correlation (Pearson and Henrici, 1896).

#### 2.2.2 Imposed stretch-shortening cycles — constant MTC shortening/lengthening velocity

To systematically explore the influence of SSC parameters on the maximal attainable AMPO, we derived predictions with a Hill-type MTC model, using estimated MTC properties specific to each rat. Specifically, we examined: 1) how the effect of MTC length excursion on the maximal attainable AMPO varied with FTS; 2) how sensitive the maximal attainable AMPO was to changes in SSC parameters and 3) how optimal values of cycle frequency, FTS, and MTC length excursion co-vary in order to maximise AMPO.

During the imposed SSC, MTC shortening and lengthening velocities were set to constant values throughout their respective phases — thus imposing no constraint on MTC acceleration — in order to explore a broad range of cycle frequency, FTS, and length excursion combinations. The maximal attainable AMPO was predicted for 2280 combinations of cycle frequency (0.5–6.0 Hz in 0.5 Hz steps), FTS (0.05–0.95 in 0.05 steps), and MTC length excursion (2–11 mm in 1 mm steps). For each combination and each rat, we optimised the muscle stimulation onset time (i.e., the time instance at which stimulation switches from 0 to its maximal value) and muscle stimulation duration (i.e., the time period during which stimulation remains at its maximal value before returning to 0) was optimised using the Nelder-Mead simplex method (Gao and Han, 2012).

To ensure periodic behaviour, optimisations were run for at least five cycles and continued thereafter until the difference in GM force at the start and end of the cycle was less than 20 mN. To verify that the global optimum had been found, we systematically varied stimulation onset time and duration. No increase in maximal attainable AMPO was observed across any SSC following changes in muscle stimulation.

Finally, as the simulations also served for a comparison with the experimental results of Askew & Marsh (1997; 1998), we reduced the SEE slack length in our simulations. In their setup, a clip was attached to the distal tendon as closely as possible to the muscle fibres, while the proximal tendon was left intact. Based on this, we performed the SSCs with a reduced SEE slack length of 3 mm for all rats.

#### 2.2.3 Imposed stretch-shortening cycles — optimal MTC shortening/lengthening velocity

The last aim in this study was to identify the MTC length over time for maximal AMPO, when MTC length over time was entirely free to take any form. This allowed us to quantify how much more AMPO could be produced when the SSC shape was completely unconstrained, compared to SSCs with constant MTC shortening and lengthening velocities. In the Hill-type MTC model used, SEE and PEE were assumed to be purely elastic (e.g., Anderson and Pandy, 1999; Soest and Bobbert, 1993) and therefore did not contribute to AMPO. Hence, only CE behaviour affected AMPO. The optimisation problem was thus reduced to finding the periodic CE length and stimulation over time in order to maximise AMPO. These optimisation problems were solved with Optimal Control techniques (see Section S3) for imposed cycle frequencies ranging from 0.5 to 6.0 Hz. Out of 615 optimisations, 604 converged successfully. Among these, the difference between maximum AMPO and the submaximal solutions was 0.3±1.1%.

## 3 Results

### 3.1 In situ experiment

#### 3.1.1 Reliability of experimental force measurements

To investigate the influence of SSC parameters on the maximal attainable AMPO of only GM, it was essential to have minimal mechanical interaction between GM and GL+PL. Our setup proved effective: GL+PL force remained below 0.2 N in all rats and conditions, with a maximum within-trial standard deviation of 15 mN. For reference, typical values of maximal isometric force of GL+PL are about 14 N (Bernabei et al., 2015), while the maximal isometric force of GM in our study was about 16 N on average. We therefore concluded that the effect of GL+PL on GM was negligible.

Several checks were used to assess the stability of GM force throughout the experiment. During the first part of the experiment (i.e., the quick-release, step-ramp, and isometric protocols), the maximum isometric GM force in the step-ramp experiments exhibited a coefficient of variation below 1%. This same low coefficient of variation (<1%) was also observed for the maximum isometric GM force during the isometric contractions in the second part of the experiment (i.e., during the SSCs). The repeated quick-release experiments revealed that GM force was ∼11 ± 8% lower midway compared to the start of the experiment, with no change midway compared to the end of the experiment. Maximal isometric GM force did, however, slightly decrease over the entire experiment. This suggests, in line with literature (e.g. Aubert et al., 1951), that the decrease in GM force was more likely caused by a shift in the MTC force-length relationship, potentially to a decrease in SEE stiffness. Based on the repeated quick-release experiment, we estimated the shift of the MTC force-length relationship to be ∼0.5 ± 0.2 mm midway compared to the start of the experiment, with no change midway compared to the end of the experiment. GM force during the ramp was ∼5 ± 3% lower midway compared to the start of the experiment, but remained stable in the second part of the experiment. With regard to the SSCs, we found that the second- and third-highest experimentally measured AMPO were only 4 ± 3% lower than the highest value across all SSC conditions and rats. Additionally, GM force development was similar between the first and second trial, which had similar muscle stimulation onset times. As intended, the second trial yielded a higher AMPO (13 ± 11% averaged over all SSC conditions and rats) due to an adjusted stimulation duration. Overall, GM behaviour was stable and reproducible during the whole experiment.

#### 3.1.2 MTC property estimation

The estimated MTC properties showed remarkable consistency across all rats (Table 1). Based on the estimated MTC properties, we derived the maximum CE shortening velocity (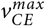 ; see Table 1), the maximum instantaneous CE power output (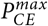 ; see Table 1) and the CE velocity at which 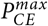 occurs 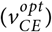 (see Table 1). Notably, the time to reach 50% of maximal isometric CE force was relatively long — approximately 34 ms at MTC optimum length in all rats. In addition to delays caused by excitation dynamics, the relatively compliant SEE substantially influenced GM force development. To exclude the influence of SEE, we calculated the time to reach 50% of maximal isometric force while holding CE length constant at its optimal value. The resulting value was considerably shorter (∼12 ms vs ∼34 ms; see Table 1) and provides a better comparison to the half-rise times at constant CE length reported in literature.

**Table 1.**
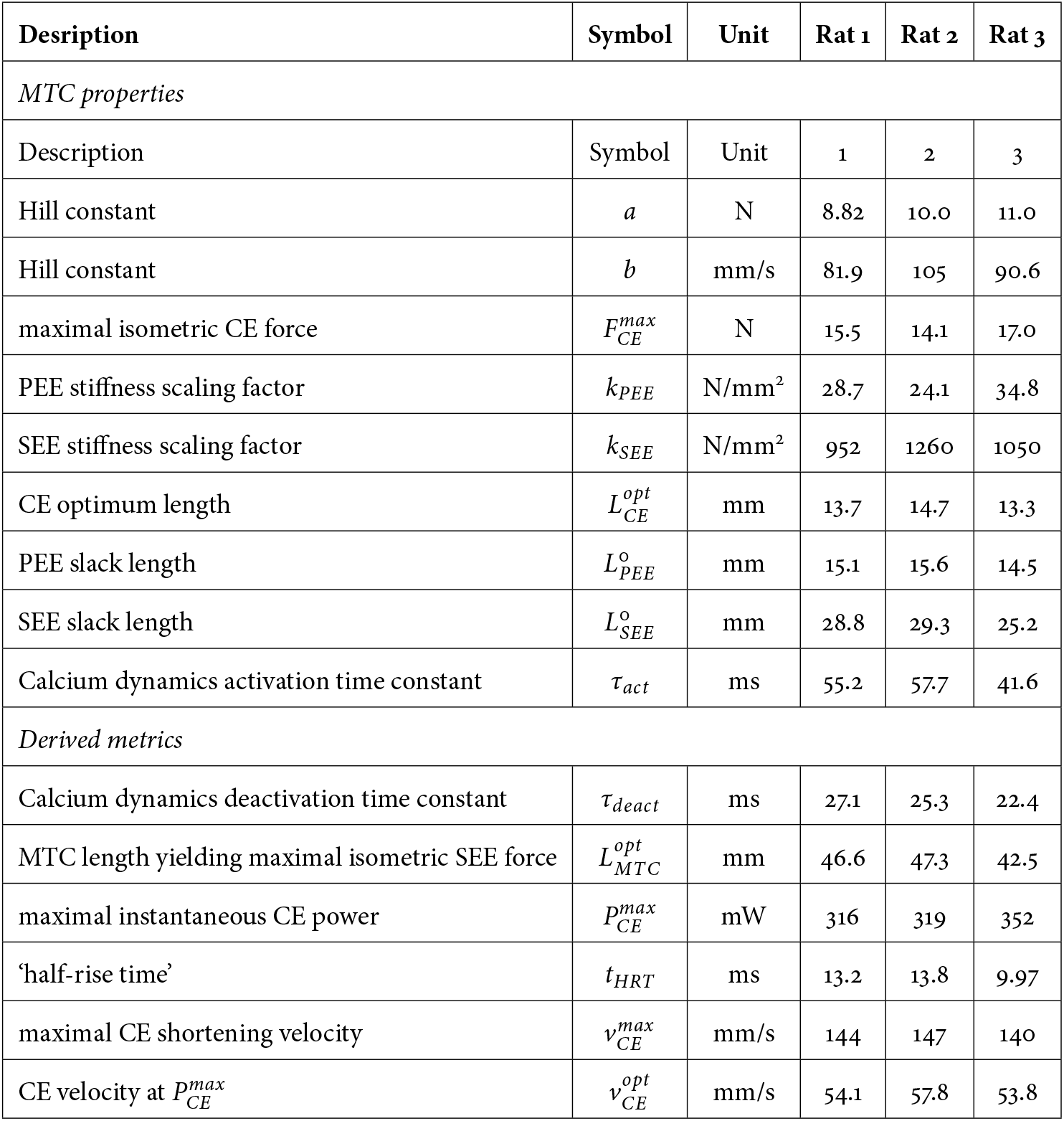
Estimated MTC properties as well as metrics derived from these properties.

| Desription | Symbol | Unit | Rat 1 | Rat 2 | Rat 3 |
| --- | --- | --- | --- | --- | --- |
| *MTC properties* |  |  |  |  |  |
| Description | Symbol | Unit | 1 | 2 | 3 |
| Hill constant | $a$ | N | 8.82 | 10.0 | 11.0 |
| Hill constant | $b$ | mm/s | 81.9 | 105 | 90.6 |
| maximal isometric CE force | $F_{CE}^{max}$ | N | 15.5 | 14.1 | 17.0 |
| PEE stiffness scaling factor | $k_{PEE}$ | N/mm<sup>2</sup> | 28.7 | 24.1 | 34.8 |
| SEE stiffness scaling factor | $k_{SEE}$ | N/mm<sup>2</sup> | 952 | 1260 | 1050 |
| CE optimum length | $L_{CE}^{opt}$ | mm | 13.7 | 14.7 | 13.3 |
| PEE slack length | $L_{PEE}^o$ | mm | 15.1 | 15.6 | 14.5 |
| SEE slack length | $L_{SEE}^o$ | mm | 28.8 | 29.3 | 25.2 |
| Calcium dynamics activation time constant | $\tau_{act}$ | ms | 55.2 | 57.7 | 41.6 |
| *Derived metrics* |  |  |  |  |  |
| Calcium dynamics deactivation time constant | $\tau_{deact}$ | ms | 27.1 | 25.3 | 22.4 |
| MTC length yielding maximal isometric SEE force | $L_{MTC}^{opt}$ | mm | 46.6 | 47.3 | 42.5 |
| maximal instantaneous CE power | $P_{CE}^{max}$ | mW | 316 | 319 | 352 |
| ‘half-rise time’ | $t_{HRT}$ | ms | 13.2 | 13.8 | 9.97 |
| maximal CE shortening velocity | $v_{CE}^{max}$ | mm/s | 144 | 147 | 140 |
| CE velocity at $P_{CE}^{max}$ | $v_{CE}^{opt}$ | mm/s | 54.1 | 57.8 | 53.8 |

#### 3.1.3 Stretch-shortening cycles

The experimentally measured AMPO of the SSC conditions are depicted in Figure 5 and detailed in Table S1 and Table S2. For brevity, we refer to a SSC condition with a cycle frequency of 2 Hz and MTC length excursion of 8 mm as 8 mm@2 Hz. In the SSC conditions with FTS set to 0.5, experimentally measured AMPO peaked for ∼4 mm@4 Hz and for ∼8 mm@2.5 Hz (Figure 5). This indicates that the cycle frequency that maximises AMPO depends on the MTC length excursion. Average MTC velocity was 32 mm/s for 4 mm@4 Hz, and 40 mm/s for 8 mm@2.5 Hz. Since GM force was almost zero at the start and end of shortening, average CE shortening equalled average MTC shortening velocity. This shows that average CE velocity that maximises AMPO should be substantially lower than the CE velocity at which instantaneous mechanical power output is maximal (∼55 mm/s; Table 1).

**Figure 5:**
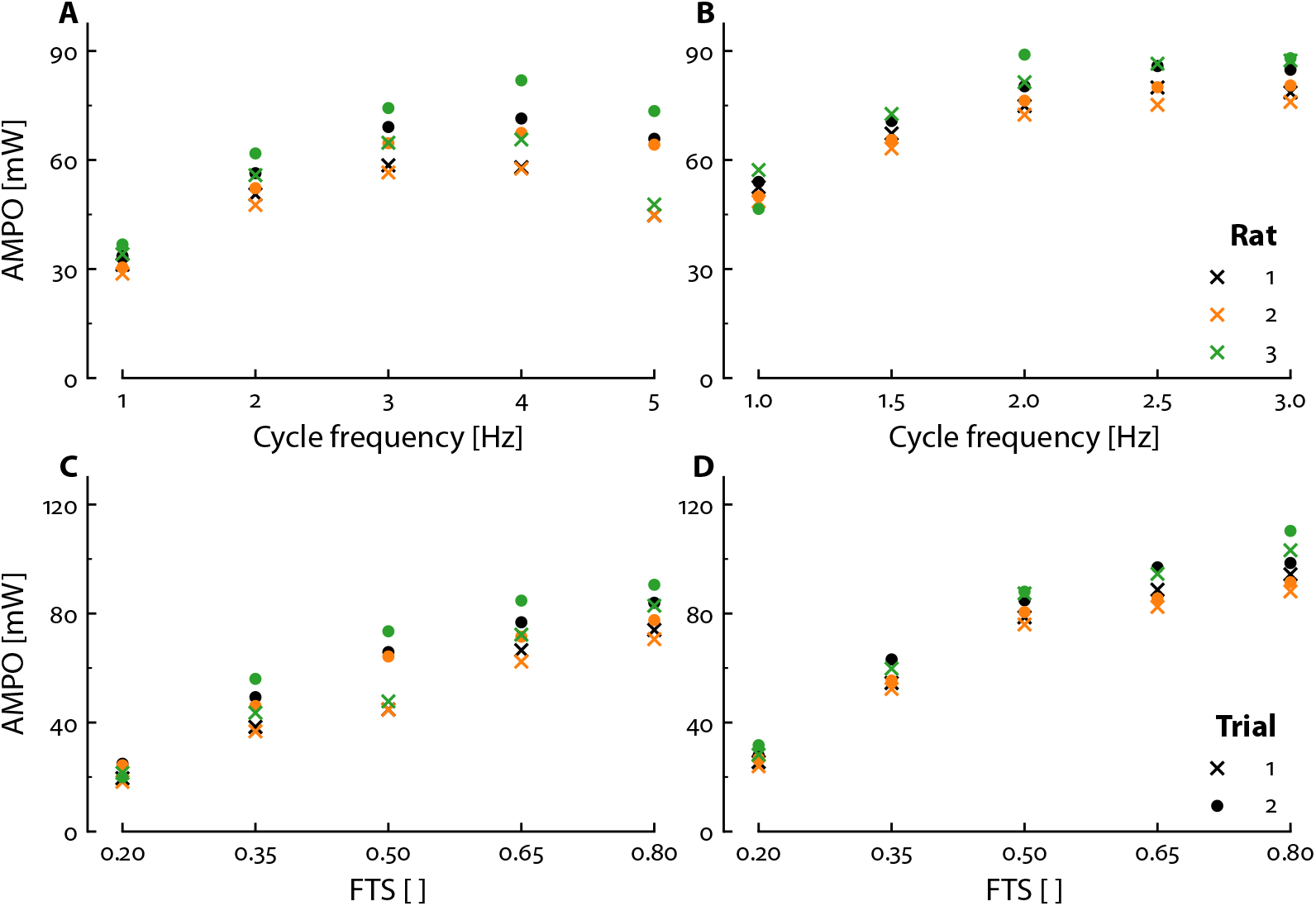
Experimentally measured influence of cycle frequency and FTS on AMPO. AMPO as a function of cycle frequency, with a fixed FTS of 0.5 at an MTC length excursion of 4 mm (A) and 8 mm (B). AMPO as a function of FTS, with a fixed cycle frequency of 3 Hz at an MTC length excursion of 4 mm (C) and 8 mm (D). Each combination of cycle frequency, FTS and MTC length excursion was performed twice. The first rial with suboptimal muscle stimulation duration and the second trial with an improved muscle stimulation duration.

In the SSC conditions with fixed cycle frequency, experimentally measured AMPO increased with FTS, reaching its highest measured value at the maximum imposed FTS of 0.8 (Figure 5). This relationship was consistent across both 4 mm and 8 mm MTC length excursions, suggesting that the FTS yielding maximal AMPO is largely independent of MTC length excursion. The effect of FTS on AMPO was most pronounced at low cycle frequencies. For example, raising FTS from 0.2 to 0.5 increased AMPO by ∼205% for 4 mm@3 Hz and ∼185% for 8 mm@2 Hz, while the increase in AMPO from FTS 0.5 to 0.8 was substantially smaller with 20% for both 4 mm@3 Hz and 8 mm@2 Hz (Figure 5; Table S1; Table S2). Since the effect of increasing FTS from 0.65 to 0.80 was small — 8% for 4 mm@3 Hz and 4% for 8 mm@2 Hz — we conclude that the FTS yielding maximal AMPO for these combinations of cycle frequency and MTC length excursion was close to 0.8. Lastly, the effect of FTS on AMPO became more pronounced at higher cycle frequencies. For example, increasing FTS from 0.5 to 0.8 increased AMPO by ∼50% for 4 mm@5 Hz and ∼40% for 8 mm@3 Hz, which is substantially more than the 8% and 4% increase for 4 mm@3 Hz and 8 mm@2 Hz (see Table S1; Table S2).

### 3.2 MTC model

#### 3.2.1 Evaluation of measured versus predicted maximal attainable AMPO

We aimed to explore the influence of SSC parameters – cycle frequency, FTS and MTC length excursion – on the maximal attainable AMPO across a broad SSC parameter space. As discussed earlier, such a comprehensive investigation is not feasible experimentally because only a limited set of conditions can be tested. Therefore, we used a Hill-type MTC model to investigate a broad SSC parameter space. For this modelling approach to be meaningful, it was crucial that the model could accurately predict the influence of MTC length and stimulation over time on AMPO.

The model-predicted GM force closely matched the measured GM force during activation across all rats and SSC conditions (Figure 6). During relaxation, measured GM force was slightly lower than model-predicted GM force in most conditions. As a result, measured AMPO was on average 6 ± 13% lower than predicted. Nonetheless, the correlations between measured and model-predicted AMPO were nearly perfect (r^2^ = 0.99, r^2^ = 0.99 and r^2^ = 0.98 for rat 1, 2 and 3 respectively; Figure 7). This was a remarkable result, as neither the model inputs nor the estimated MTC properties were tuned to experimentally measured SSCs. These findings demonstrate that a Hill-type MTC model can accurately predict the influence of muscle length and stimulation over time on AMPO.

**Figure 6:**
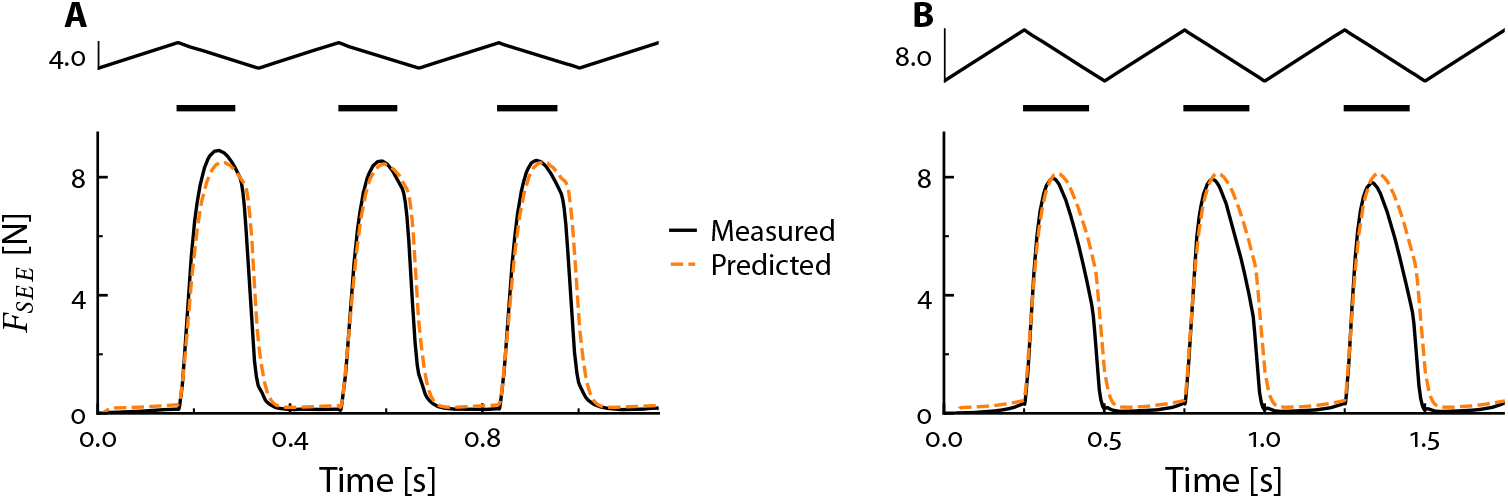
Comparison of experimentally measured and predicted GM force over time. Predicted GM force over time was derived using a Hill MTC-type model, with experimentally measured MTC length and stimulation over time as inputs. A) SSC with a cycle frequency of 3 Hz, a FTS of 0.5 and an MTC length excursion of 4 mm. B) SSC with a cycle frequency of 2 Hz, a FTS of 0.5 and an MTC length excursion of 8 mm.

**Figure 7:**
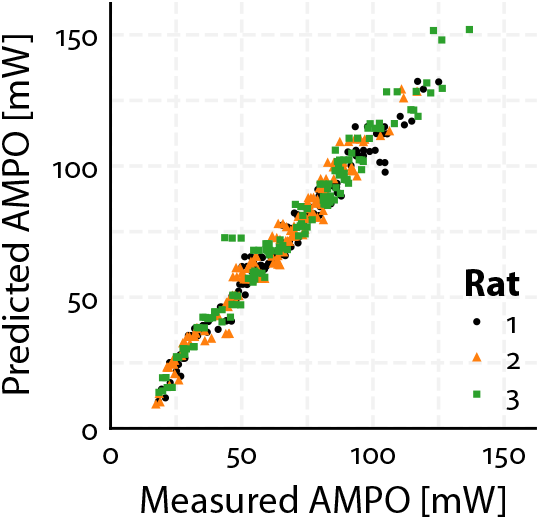
Comparison of experimentally measured and predicted AMPO. Measured AMPO was derived from experimentally observed MTC length and GM force. Predicted AMPO was derived using a Hill MTC-type model, with experimentally measured MTC length and stimulation over time as inputs to predict GM force. Each dot represent the AMPO of one full cycle where muscle stimulation was present, such that there are three dots for each experimental SSC condition (see Section 2.1.6). The nearly perfect correlation demonstrates that a Hill-type MTC model can accurately predict the influence of MTC length and stimulation over time on AMPO.

#### 3.2.2 Imposed stretch-shortening cycles — constant MTC shortening/lengthening velocity

Using the Hill-type MTC model, we predicated that AMPO peaks at a value of 169 mW (averaged across the three rats) at cycle frequency of 3.5 Hz, an FTS of 0.85 and an MTC length excursion of 8 mm (Figure 8 & Figure 9). The optimal FTS remained remarkably constant across all tested combinations of cycle frequency and MTC length excursion (Figure 9). By contrast, cycle frequency and MTC length excursion showed a strong interaction, such that when cycle frequency was increased, the optimal MTC length excursion decreased, and vice versa (Figure 9). Notably, a broad range of combinations of SSC parameters achieved AMPO values close to its peak value (Figure 8 and Figure 9). For instance, cycle frequency ranging between 2.9 and 4.5 Hz produced an AMPO within 95% of maximum AMPO, when adjusting FTS and MTC length excursion accordingly. For FTS this was a range between 0.74 and 0.91, when adjusting cycle frequency and MTC length excursion accordingly, while it was between 6.2 and 10.2 mm for the MTC length excursion when adjusting cycle frequency and FTS accordingly. These findings demonstrate that while a clear optimum exists for maximal AMPO, there is a wide variety of combinations of SSC parameters that yield near-maximal AMPO values.

**Figure 8:**
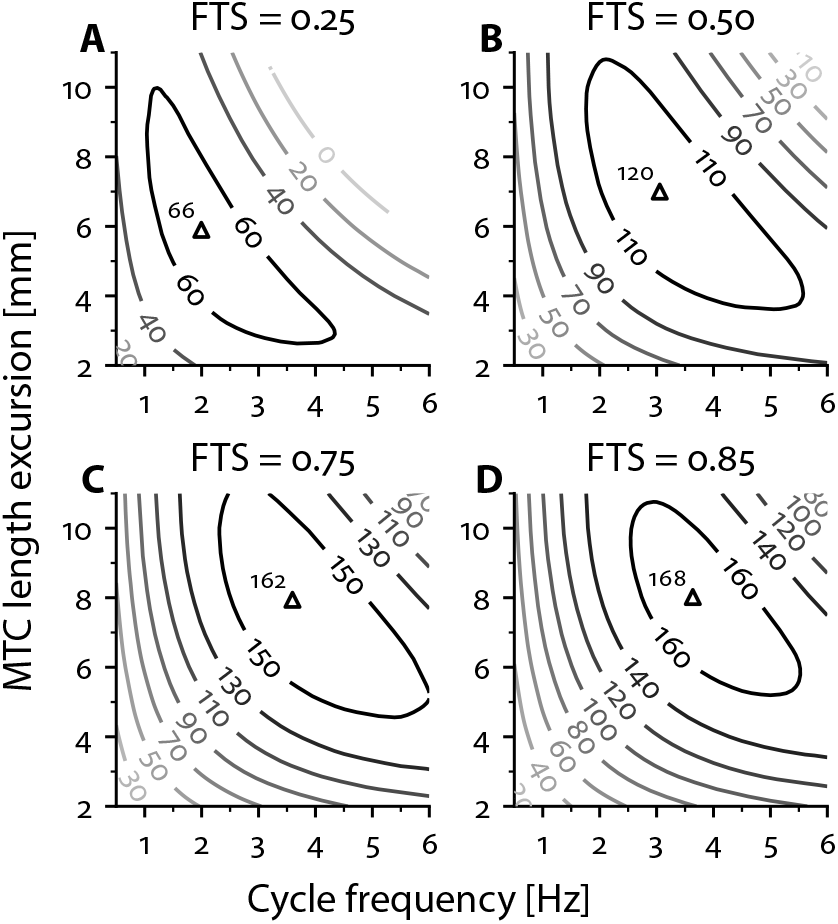
Predicted maximal attainable AMPO as a function of cycle frequency and MTC length excursion, shown for four distinct FTS values. The maximal attainable AMPO (in mW), averaged across three rats, are depicted as contour lines. The open triangles indicate the location of peak AMPO at each FTS corresponding to the optimal combination of cycle frequency and MTC length excursion for each FTS, with the corresponding peak AMPO labelled at the top-left of the triangle.

**Figure 9:**
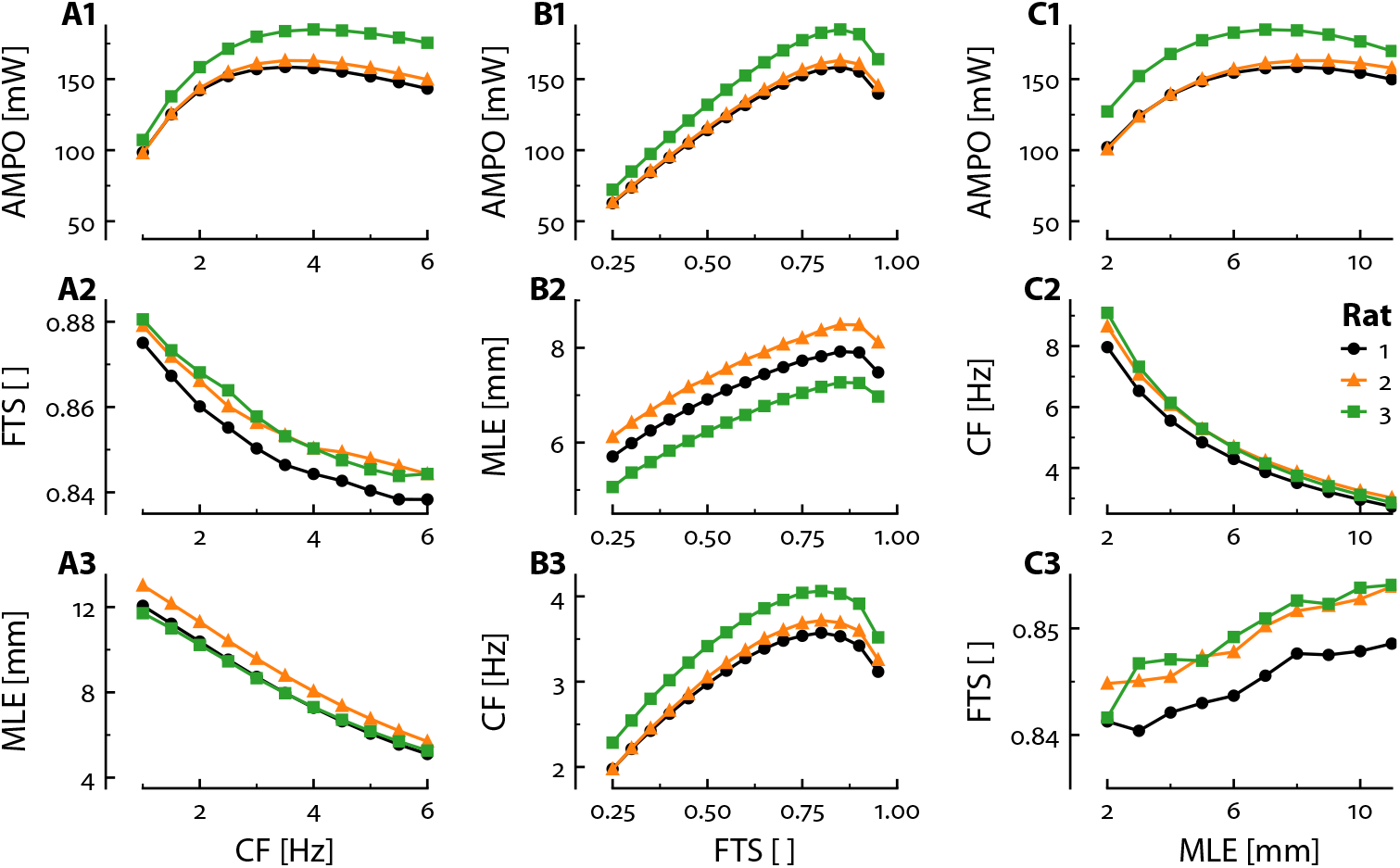
Predicted interrelationships between cycle frequency (CF), FTS, MTC length excursion (MLE) and the maximal attainable AMPO. These plots collectively illustrate how cycle frequency, FTS, and MTC length excursion interact to maximise AMPO under different conditions. This figure presents a 3×3 grid of plots created by imposing one SSC parameter at a time while optimising the two other SSC parameters to maximise AMPO. In the first column, cycle frequency was imposed, and AMPO (A1) was maximised by identifying the optimal FTS (A2) and MTC length excursion (A3) at each cycle frequency. In the second column, FTS was imposed, and AMPO (A1) was maximised by identifying the optimal MTC length excursion (B2) and cycle frequency (B3) at FTS. In the third column, MTC length excursion was imposed, and AMPO (C1) was maximised by identifying the optimal cycle frequency (C2) and FTS (C3) at each MTC length excursion. For all columns, muscle stimulation onset time and duration were optimised to maximise AMPO.

#### 3.2.3 Imposed stretch-shortening cycles — optimal MTC shortening/lengthening velocity

Using the Hill-type MTC model, we identified the optimal CE length over time for maximal AMPO without imposing any constraints MTC length over time (in contrast to section Section 3.2.2, where MTC shortening and lengthening velocity was constant). We found that AMPO was maximised when CE shortened at an approximately constant velocity during the shortening phase and lengthened with increasing velocity during the lengthening phase (see Figure 10), across all imposed cycle frequencies. Given CE length and CE force over time (and PEE and SEE properties), the corresponding MTC length over time can be calculated. Thus, MTC length over time is inherently dependent on the properties of SEE. Because SEE force is not constant during CE shortening, the MTC velocity that maximises AMPO is generally not constant during the shortening phase; even if CE shortening velocity is constant. The size of this effect depends on the SEE slack length: with a typical slack length (e.g., 28 mm), the MTC velocity is clearly not constant (see Figure 10E), whereas with a short slack length (e.g., 3 mm), the MTC velocity closely resembles the CE velocity and is therefore nearly constant during shortening (see Figure 10F).

**Figure 10:**
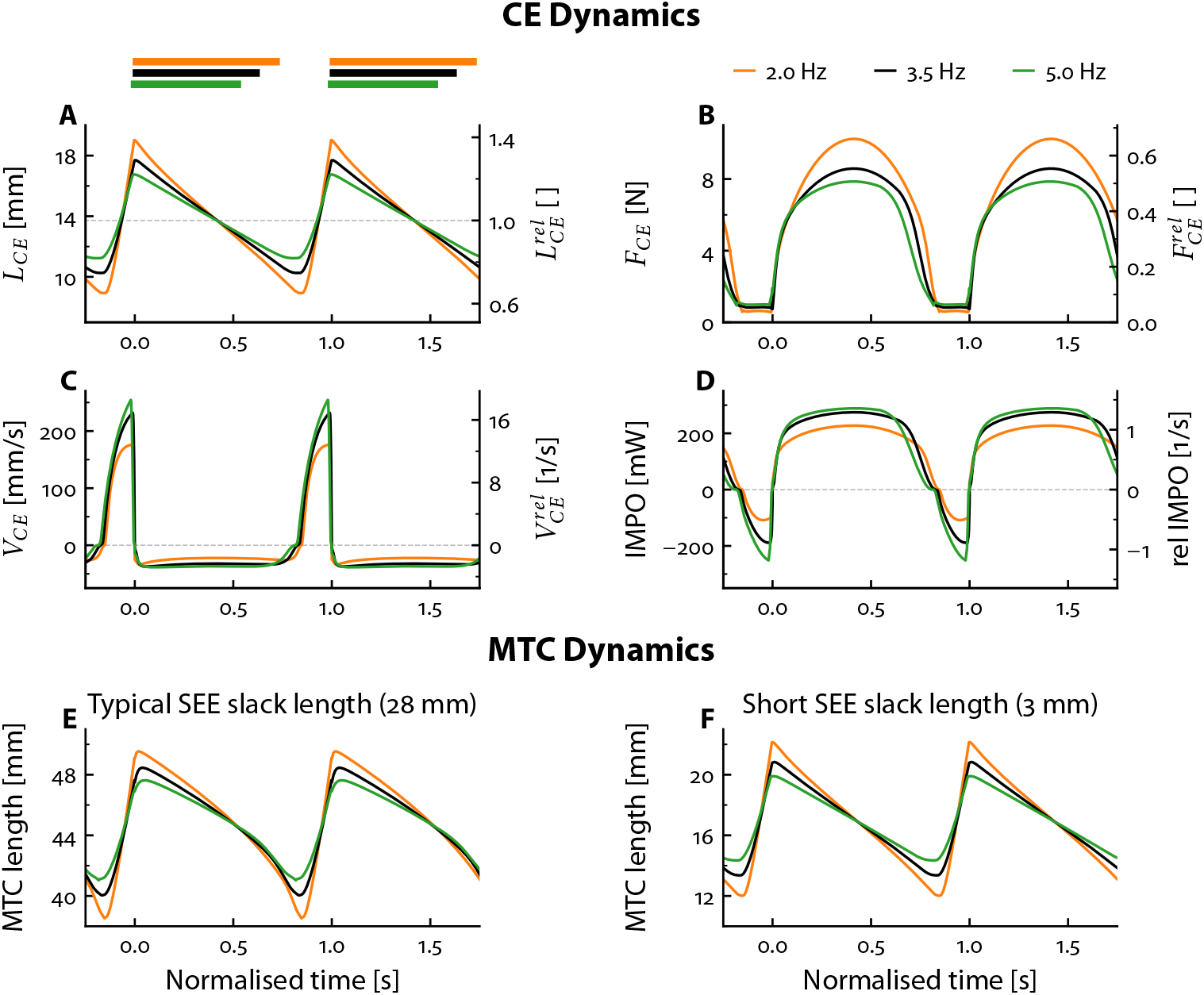
Predicted CE behaviour for maximal attainable AMPO, shown at three distinct cycle frequencies for rat 1. CE length (A), CE force (B), CE velocity (C) and instantaneous mechanical power output (IMPO) (D) as a function of normalised time (i.e. time divided by the cycle duration). CE stimulation was maximal during the period indicated by the coloured bars and fully off elsewhere. The right labels depict the normalised values, where CE length and CE velocity are normalised to optimum CE length, CE force is normalised to maximal isometric CE force and IMPO is normalised to the product of optimum CE length and maximal isometric CE force. Based on CE length and force over time, and the SEE properties, the MTC length over time can be calculated. The resulting MTC length over time substantially differs between a typical SEE slack length of 28mm (E) and a short SEE slack length of 3 mm (F).

#### 3.2.3 CE Dynamics

For a short SEE slack length, the MTC length over time that maximises AMPO has a near-constant shortening and lengthening velocity. As a result, the maximum attainable AMPO using the optimised CE length over time was only 2 ± 1% higher than that achieved with the constant MTC shortening and lengthening velocities with a short SEE slack length. It should be noted, however, this small increase in AMPO is specific to imposed cycle frequency. When imposing FTS or MTC length excursions, the optimal CE length over time was substantially different than that with constant MTC velocities during shortening and lengthening – especially at suboptimal values. For the interested reader, we refer to the Supplementary Material (Section S4) for the effect of FTS or MTC length excursion on the optimal MTC behaviour for maximising AMPO.

## 4 Discussion

The aim in this study was to explore the influence of SSC parameters — cycle frequency, FTS and MTC length excursion — on the maximal attainable AMPO of rat GM. To this end, we combined *in situ* experiments with simulations and optimisations using a Hill-type MTC model. Model parameter values were estimated individually for each specimen based on independent data from quick-release, step-ramp and isometric experiments. Additionally, the model inputs (i.e., MTC length and stimulation over time) were directly derived from the experimental SSC data. The correlation coefficients between model-predicted and measured AMPO were nearly perfect (r^2^ > 0.98). This is a remarkable result, particularly given that the predictions were not tuned to any of the experimental SSCs. This strong correspondence enabled us to leverage both experimental findings and model-based predictions to investigate how SSC parameters influence the maximal attainable AMPO.

Both experimental data and model-based predictions showed that the MTC should spend substantially more time shortening than lengthening in order to maximise AMPO — at an FTS of approximately 0.85. This optimal FTS value remained remarkably constant across all tested combinations of cycle frequency and MTC length excursion. By contrast, cycle frequency and MTC length excursion showed a strong interaction, such that when cycle frequency increased, the optimal MTC length excursion decreased, and vice versa. Lastly, we found that AMPO peaked at a cycle frequency of 3.5 Hz, an FTS of 0.85 and at an MTC length excursion of 8 mm, with a relatively constant CE velocity throughout the shortening phase and an increasing CE lengthening velocity throughout the lengthening phase.

### 4.1 The optimal FTS for AMPO was relatively constant across all SSC conditions

Our results show that the maximal attainable AMPO substantially increased when FTS increased from 0.25 to 0.50, regardless of cycle frequency and/or MTC length excursion. For example, at the optimal combination of cycle frequency and MTC length excursion, increasing FTS from 0.25 to 0.50 caused AMPO to increase by about 95%. This finding is consistent with the results of Askew and Marsh (1997), who reported increases of ∼90% for m. soleus and ∼70% for m. extenstor digitorum longus when FTS increased from 0.25 to 0.50. Further increasing FTS from 0.50 to 0.75 yielded an additional 30% increase in AMPO, aligning with Askew & Marsh’s observations of a ∼40% increase in AMPO for both muscles. Moreover, Askew and Marsh (1997) observed that AMPO increased even further, and peaked at an FTS between 0.80 and 0.90 at the optimal combination of cycle frequency and MTC length excursion. Our results not only corroborate this finding, but also demonstrate that the optimal FTS was relatively constant (ranging 0.84–0.88) across all tested cycle frequencies (1–6 Hz) and MTC length excursions (1-11 mm).

The optimal FTS reflects mainly a trade-off between the CE concentric force–velocity relationship and excitation dynamics. The negative effects of both excitation dynamics and the force–velocity relationship on net mechanical work per cycle increase with cycle frequency. Yet, the fact that the optimal FTS remains relatively constant across all cycle frequencies studied suggests that these negative effects increase to a similar extent with cycle frequency (Figure 9).

### 4.2 The effect of FTS on optimal cycle frequency and MTC length excursion for maximising AMPO

While the optimal FTS remained relatively constant across all cycle frequencies and MTC length excursions, FTS itself strongly influenced the optimal values of both cycle frequency and MTC length excursion (Figure 9). With respect to cycle frequency, we observed that the optimal value increased with FTS, although the magnitude of this increase diminished at higher FTS values. This observation aligns well with findings by Askew and Marsh (1997), who reported comparable increases across FTS values for both m. soleus and m. extensor digitorum longus. For MTC length excursion, we found that the optimal value increased with 25% and 10% when raising FTS from 0.25 to 0.50 and from 0.50 to 0.75, respectively. A direct comparison with Askew and Marsh (1997) is difficult, as their results differed between muscles: for m. soleus, these increases were 35% and 40%, while they were 0% and 35% for m. extensor digitorum longus. Lastly, Askew and Marsh (1997) also examined how average muscle shortening velocity varied with FTS. For m. extensor digitorum longus, they found a consistent decrease in optimal shortening velocity as FTS increased. By contrast, m. soleus showed a slight increase in optimal shortening velocity, although this discrepancy may stem from missing data near the optimal frequency at FTS = 0.75 (see their Figure 4). A follow-up study by the same authors (Askew and Marsh, 1998) later confirmed that the optimal average shortening velocity for m. soleus indeed decreases with increasing FTS, independently of cycle frequency. We observed the same for our model-predictions. Overall, our findings regarding the influence of FTS on optimal cycle frequency, MTC length excursion, and average shortening velocity are in line with previous literature. These findings are not surprising, given the effect of the CE force-velocity relationship. After all, at higher FTS, there is relatively more time for shortening than at lower FTS values. This enables an increase in cycle frequency and allows muscles to shorten and lengthen over a greater distance, without substantial negative effects due to the effect of shortening velocity on muscle force.

### 4.3 Optimal cycle frequency and optimal MTC length excursion for maximal AMPO

While the optimal FTS for maximal AMPO remained relatively constant across all SSCs studied, we found that cycle frequency and MTC length excursion strongly interacted for maximal AMPO. Specifically, as cycle frequency increased, the optimal MTC length excursion for the maximal attainable AMPO decreased, and vice versa (see Figure 8 & Figure 9). This interaction has previously been observed in sinusoidal SSCs (e.g., James et al., 1996) – where shortening and lengthening durations are equal. However, to our knowledge, it has not been reported for SSCs with unequal shortening and lengthening durations; although Askew and Marsh (1997) briefly noted this interaction, they did not present the relevant data.

The interaction between cycle frequency and MTC length excursion for maximal attainable AMPO did not come as a surprise, given the negative effect of the force-velocity relationship on AMPO. In line with Askew and Marsh (1997), Caiozzo and Baldwin (1997) and James et al. (1996), we observed that the maximal attainable AMPO was less sensitive to increases in cycle frequency and MTC length excursion above their respective optimal values than to decreases below these optimal values. This is not only true at optimal FTS (see Figure 9), but also for suboptimal FTS (see Figure 8, where contour lines below the optimum cycle frequency/MTC length excursion are closer to their optimum value than values above their optimum). The finding that AMPO was less sensitive to increases in cycle frequency and MTC length excursion above their respective optimal values than to decreases below these optimal values might be counter-intuitive, as one may expect the maximal attainable AMPO to decline rapidly due a higher shortening velocity (at higher cycle frequencies and MTC length excursion). However, at low cycle frequencies, the MTC must shorten and lengthen over a substantial distance, which reduces the maximal attainable AMPO primarily due the effect of the CE force-length relationship. Conversely, at low MTC length excursions, very high cycle frequencies are required to achieve substantial power output, which in turn compromises AMPO due to the effect of the CE force-velocity relationship (i.e., muscle contracts at lengths at which CE force is substantially lower than the maximal isometric force). In sum, our findings indicate that deviations above the optimal cycle frequency and MTC length excursion have a less detrimental effect on the maximal attainable AMPO than deviations below them.

### 4.4 The effect of SEE properties on the MTC behaviour for maximal AMPO

To maximise AMPO, CE should shorten at an approximately constant velocity during the shortening phase and lengthen with increasing velocity during the lengthening phase (see Figure 10). The resulting MTC length over time for maximal AMPO is determined not only by CE length over time, but also by the SEE properties — particularly SEE compliance and SEE slack length. For typical SEE properties, MTC velocity should be higher (i.e., less negative or even positive) than that of CE during force development, and lower (i.e., more negative) during force relaxation to maximise AMPO (Figure 10E).

The same principle applies the other way around: when velocity of MTC is constant, that of CE varies during force development and relaxation. Specifically, changes in CE velocity increase with increasing SEE compliance and SEE slack length. In the experimentally observed SSCs, the estimated SEE slack length was substantially longer (∼28 mm) than that used in the simulated SSCs (3 mm), resulting in more variable CE velocity during shortening. This variable CE velocity compromises AMPO: for a given shortening distance, CE should ideally shorten at a constant velocity to maximise AMPO. This more variable CE velocity during the shortening phase of the experimental SSC was the main reason why AMPO in the experimental SSCs was lower than in the simulated SSCs. For instance, at a 2 Hz cycle frequency, FTS of 0.5 and an 8 mm MTC length excursion, AMPO in the simulated SSC was ∼40% higher (∼80 mW vs. ∼110 mW; Figure 5 vs. Figure 8).

We found that – especially at a low FTS, a high cycle frequency and a small MTC length excursions – muscle stimulation should occur before MTC shortening starts in order to maximise AMPO. Consequently, CE shortening started before MTC shortening, such that CE was shortening over a longer time interval resulting in a lower average CE shortening velocity than MTC. Thus, the behaviour of SEE does not only cause a discrepancy between instantaneous CE and MTC velocity, it also affects their average values.

Askew and Marsh (1998) reported that at low FTS values and high cycle frequencies, the average MTC shortening velocity exceeded the CE velocity that maximises instantaneous power output based on the force-velocity relationship 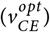. They acknowledged that this was a unexpected finding, since CE shortening velocity should theoretically be at or below 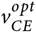 in order to maximise AMPO. Similar to our SSCs with a constant MTC shortening and lengthening velocity, Askew and Marsh (1998) left a small portion of the tendon attached to the muscle fibres. Since Askew & Marsh reported that muscle stimulation onset occurred before MTC lengthening, it is evident that average CE shortening velocity was lower than that of MTC. Therefore, the finding that optimal CE shortening velocity should exceeds 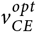 can, at least, partly be attributed to an overestimation of CE shortening velocity due to the series elastic structures.

The discrepancy between CE and MTC shortening velocity is most pronounced at low FTS, high cycle frequencies, and small MTC length excursion — exactly where Askew and Marsh (1998) observed CE shortening velocities exceeding 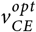. At high FTS values, however, this discrepancy is lower. Askew and Marsh (1998) found that at high FTS values (i.e., 0.75), the optimal average shortening velocity was consistently below 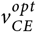. This aligns with our findings, even when muscle stimulation onset preceded MTC shortening. In conclusion, while other factors may play a role, we are confident that such factors have little influence on the effect of SSC parameters on the maximal attainable AMPO near the optimum.

### 4.5 Applications of findings

Our results reinforce the findings by Askew and Marsh (1997) that muscles should spend substantially more time shortening than lengthening in order to maximise AMPO. Obviously, many animals can only minimally influence the shortening versus lengthening duration of their muscles due to their intricate mechanical interaction with the environment. For example, during running, leg extensor muscles typically spend more time lengthening than shortening, as ground contact time is much shorter than flight time. In fact, longer shortening than lengthening durations have been observed only for m. pectoralis major in flying birds (e.g., Askew and Marsh, 2001; Biewener et al., 1998; Earls, 2000; Ellerby and Askew, 2007; Jackson and Dial, 2011; Spedding, 1987; Williamson et al., 2001) and for m. obliquus externus in calling frogs (Girgenrath and Marsh, 1997).

Human, however, can substantially influence the shortening versus lengthening duration of their muscles by using external equipment that (partially) imposes (muscle) kinematics. A prime example is the bicycle, where the cranks are mechanically coupled, causing the legs to move in anti-phase which means that the downstroke duration of one leg equals the upstroke duration of the other. As a result, muscle shortening and lengthening durations are more or less equal (FTS ≈ 0.5). Obviously, in sprint cycling, the goal is not to maximise AMPO of a single muscle, but rather to maximise the sum of AMPOs of all muscles involved, which requires an additional trade-off. Nevertheless, it stands to reason that larger muscles should operate closer to their individual optimum than smaller muscles in order to maximise the sum of AMPOs. In this regard, it is noteworthy that in single-leg cycling, the maximal attainable AMPO summed across all muscles involved can be increased by 4% when the downstroke duration was increased to 58% of the cycle duration (Martin et al., 2002). Based on our findings, we believe that such redesigns of equipment – which allows the larger muscles to realise a longer shortening than lengthening duration – can greatly improve the maximal attainable AMPO during tasks such as wheelchair riding, cycling and rowing.

## Acknowledgments

The authors thank Guus Baan for creating the illustration used in Figure 3.

## Data and resource availability

All data, code, and materials used in this study are openly available:

- **GitHub repository:** All raw data, processed data, and analysis code are hosted on GitHub at https://github.com/edwinreuvers/rat-gm-ampo.
- **Reproducible analysis website:** Full analysis pipeline — including data, analysis code, and figure/table generation — is available at https://edwinreuvers.github.io/publications/rat-gm-ampo.

## Supplementary material

### S1 Selection of experimental SSC conditions

In the experiment, we aimed to measure an informative and representative subset of stretch-shortening cycles (SSCs) to investigate the effect of the SSC parameters — cycle frequency, FTS and MTC length excursion — on the maximal attainable AMPO. Yet, a key question was: Which SSCs should be selected? To make an informed decision, we performed preliminary simulations using a Hill-type MTC model with parameter values from literature for three rat m. gastrocnemius medialis (Zandwijk et al., 1996).

We predicted maximal AMPO across various combinations of cycle frequency and FTS at five distinct MTC length excursions (2, 4, 6, 8 and 10 mm), each centred at an average MTC length ∼3 mm below the MTC length yielding maximal isometric GM force. Muscle stimulation onset time (i.e., the instance at which stimulation switches from 0 to its maximal value) was set to the start of MTC shortening. Muscle stimulation duration (i.e., the time period during which stimulation remains at its maximal value before returning to 0) was optimized using the Nelder-Mead simplex method (Gao and Han, 2012). To ensure periodic behaviour, optimisations were run for at least five cycles and until the muscle force at the beginning of the cycle deviated less than 20 mN from its value at the end of the cycle. Maximal AMPO was calculated for last cycle.

We then made contour plots for every MTC length excursion (Figure S1), showing the influence of cycle frequency and FTS on maximal AMPO at each MTC length excursion. For the experimental SSC conditions, we limited the range of FTS between 0.2 and 0.8 because values outside this range will yield very short shortening or lengthening durations. Within this FTS range, we observed that AMPO peaked at about 5 and 3 Hz for an MTC length excursion of 4 and 8 mm respectively. Again, to prevent short shortening and lengthening duration, we set the maximum cycle frequencies accordingly. Based on these findings, we selected three subsets of SSC conditions: 1) constant FTS, varying cycle frequency; 2) constant cycle frequency, varying FTS; 3) varying both cycle frequency and FTS. Because there was overlap in the SSC conditions, this yielded 13 unique SSC conditions for each MTC length excursion, such that there were 26 different SSC conditions in total.

**Figure S1:**
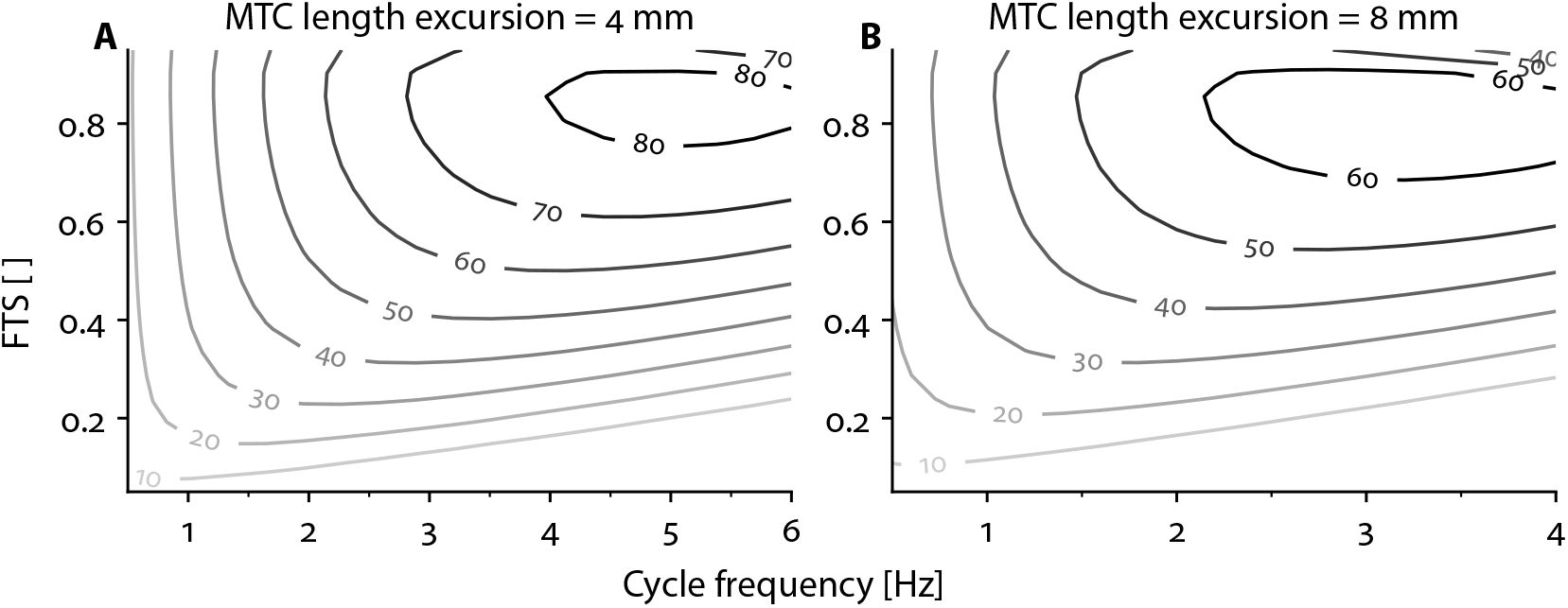
Preliminary predictions of maximal AMPO as a function of cycle frequency and FTS. The contour lines depict the maximal attainable AMPO (in mW) averaged over three parameter sets. These preliminary predictions were derived with a Hill-type MTC model with parameters obtained from literature (Zandwijk et al., 1996) and were used to inform the selection of experimental SSC conditions.

### S2 Experimental SSC conditions and experimentally measured AMPO

**Table S1:** SSC parameters, stimulation durations, and measured AMPO of experimental stretch-shortening cycles with a 4 mm MTC length excursion. Stimulation onset was set at the start of MTC shortening in all conditions.

|  |  | Condition |  |  |  |  |  |  |  |  |  |  |  |  |
| --- | --- | --- | --- | --- | --- | --- | --- | --- | --- | --- | --- | --- | --- | --- |
|  | Unit | 1 | 2 | 3 | 4 | 5 | 6 | 7 | 8 | 9 | 10 | 11 | 12 | 13 |
| SSC parameters |  |  |  |  |  |  |  |  |  |  |  |  |  |  |
| Cycle frequency | Hz | 1.0 | 2.0 | 3.0 | 4.0 | 5.0 | 3.0 | 3.0 | 3.0 | 3.0 | 5.0 | 4.0 | 2.0 | 1.0 |
| FTS | - | 0.50 | 0.50 | 0.50 | 0.50 | 0.50 | 0.80 | 0.65 | 0.35 | 0.20 | 0.80 | 0.65 | 0.35 | 0.20 |
| MTC shortening and lengthening times |  |  |  |  |  |  |  |  |  |  |  |  |  |  |
| MTC shortening time | ms | 500 | 250 | 167 | 125 | 100 | 267 | 217 | 117 | 66.7 | 160 | 162 | 175 | 200 |
| MTC lengthening time | ms | 500 | 250 | 167 | 125 | 100 | 66.7 | 117 | 217 | 267 | 40.0 | 87.5 | 325 | 800 |
| Stimulation durations |  |  |  |  |  |  |  |  |  |  |  |  |  |  |
| Trial 1 | ms | 410 | 170 | 100 | 65 | 35 | 180 | 130 | 53 | 23 | 85 | 95 | 120 | 140 |
| Trial 2 | ms | 450<sup>1</sup> | 210 | 120 | 85 | 55 | 210 | 160 | 73 | 33<sup>2</sup> | 110 | 110 | 130 | 160<sup>3</sup> |
| Measured AMPO of Rat 1 |  |  |  |  |  |  |  |  |  |  |  |  |  |  |
| Trial 1 | mW | 31 | 51 | 59 | 58 | 45 | 74 | 67 | 38 | 20 | 82 | 74 | 43 | 23 |
| Trial 2 | mW | 34 | 56 | 69 | 72 | 66 | 84 | 77 | 49 | 25 | 100 | 87 | 47 | 27 |
| Measured AMPO of Rat 2 |  |  |  |  |  |  |  |  |  |  |  |  |  |  |
| Trial 1 | mW | 29 | 48 | 57 | 58 | 45 | 71 | 62 | 37 | 18 | 79 | 70 | 40 | 22 |
| Trial 2 | mW | 30 | 52 | 65 | 68 | 64 | 78 | 72 | 46 | 24 | 93 | 79 | 45 | 24 |
| Measured AMPO of Rat 3 |  |  |  |  |  |  |  |  |  |  |  |  |  |  |
| Trial 1 | mW | 34 | 56 | 65 | 66 | 48 | 83 | 72 | 44 | 22 | 91 | 82 | 47 | 25 |
| Trial 2 | mW | 37 | 62 | 74 | 82 | 74 | 91 | 85 | 56 | 20 | 120 | 96 | 54 | 28 |

<sup>1</sup> For condition 1: stimulation duration of rat 3 was 455 ms.
<sup>2</sup> For condition 9: stimulation duration of rat 3 was 23 ms.
<sup>3</sup> For condition 13: stimulation duration of rat 1 was 165 ms.

**Table S2:** SSC parameters, stimulation durations, and measured AMPO of experimental stretch-shortening cycles with a 4 mm MTC length excursion. Stimulation onset was set at the start of MTC shortening in all conditions.

|  |  | Condition |  |  |  |  |  |  |  |  |  |  |  |  |
| --- | --- | --- | --- | --- | --- | --- | --- | --- | --- | --- | --- | --- | --- | --- |
|  | Unit | 1 | 2 | 3 | 4 | 5 | 6 | 7 | 8 | 9 | 10 | 11 | 12 | 13 |
| SSC parameters |  |  |  |  |  |  |  |  |  |  |  |  |  |  |
| Cycle frequency | Hz | 1.0 | 1.5 | 2.0 | 2.5 | 3.0 | 2.0 | 2.0 | 2.0 | 2.0 | 3.0 | 2.5 | 1.5 | 1.0 |
| FTS | - | 0.50 | 0.50 | 0.50 | 0.50 | 0.50 | 0.80 | 0.65 | 0.35 | 0.20 | 0.80 | 0.65 | 0.35 | 0.20 |
| MTC shortening and lengthening times |  |  |  |  |  |  |  |  |  |  |  |  |  |  |
| MTC shortening time | ms | 500 | 333 | 250 | 200 | 167 | 400 | 325 | 175 | 100 | 267 | 260 | 233 | 200 |
| MTC lengthening time | ms | 500 | 333 | 250 | 200 | 167 | 100. | 175 | 325 | 400 | 66.7 | 140 | 433 | 800 |
| Stimulation durations |  |  |  |  |  |  |  |  |  |  |  |  |  |  |
| Trial 1 | ms | 410 | 250 | 180 | 140 | 100 | 300 | 250 | 110 | 55 | 170 | 180 | 160 | 150 |
| Trial 2 | ms | 460 | 290 | 210 | 160 | 120 | 340 | 270 | 140 | 75<sup>1</sup> | 210<sup>2</sup> | 220 | 190 | 170<sup>3</sup> |
| Measured AMPO of Rat 1 |  |  |  |  |  |  |  |  |  |  |  |  |  |  |
| Trial 1 | mW | 53 | 67 | 75 | 80 | 79 | 94 | 89 | 55 | 26 | 110 | 95 | 54 | 34 |
| Trial 2 | mW | 54 | 71 | 80 | 86 | 85 | 99 | 97 | 63 | 28 | 120 | 100 | 62 | 35 |
| Measured AMPO of Rat 2 |  |  |  |  |  |  |  |  |  |  |  |  |  |  |
| Trial 1 | mW | 49 | 63 | 73 | 75 | 76 | 88 | 83 | 52 | 24 | 100 | 88 | 50 | 32 |
| Trial 2 | mW | 50 | 66 | 76 | 80 | 81 | 92 | 86 | 55 | 27 | 110 | 95 | 54 | 33 |
| Measured AMPO of Rat 3 |  |  |  |  |  |  |  |  |  |  |  |  |  |  |
| Trial 1 | mW | 57 | 73 | 81 | 87 | 88 | 100 | 95 | 60 | 28 | 120 | 100 | 60 | 37 |
| Trial 2 | mW | 47 | - | 89 | - | 88 | 110 | - | - | 32 | 130 | - | - | 40 |

<sup>1</sup> For condition 9: stimulation duration of rat 3 was 65 ms.<sup>2</sup> For condition 10: stimulation duration of rat 3 was 223 ms.<sup>3</sup> For condition 13: stimulation duration of rat 1 was 175 ms.

### S3 Optimisation protocol to identify optimal CE length over time

One of the aims of this study was to identify the MTC length over time for maximal AMPO, when MTC length over time is entirely free to take any form. To predict this, we used a Hill-type MTC model. In our Hill-type MTC model used, we assumed that both SEE and PEE were purely elastic elements (e.g., Anderson and Pandy, 1999; Soest and Bobbert, 1993). Therefore, neither PEE nor SEE contributed to AMPO of the MTC during fully periodic contractions. Hence, only CE behaviour influenced AMPO and therefore only CE was modelled to simplify the optimisation problem.

The optimisation problem was thus to find the periodic CE length and *STIM* over time for maximal AMPO. Optimisations were performed for 1) imposed cycle frequencies (1.0–6.0 Hz in 0.5 Hz steps); 2) imposed FTS values (0.05–0.95 in 0.05 steps) and 3) imposed MTC length excursions (2–11 mm in 1 mm steps). For all optimisations, the following cost-function was minimised:

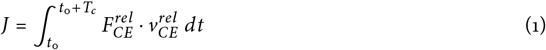

*t*_0_ denotes the initial time and *T*_*c*_ denotes the cycle time.

The solution had to satisfy the differential equation describing the activation dynamics:

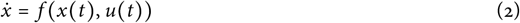

*x*(*t*) denotes that states of the system 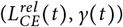 and *u*(*t*) denotes the inputs of the system 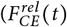 and *STIM*(*t*)).

The following task constraint was added to achieve periodic behaviour:

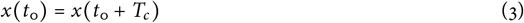

Moreover, inequality constraints were imposed on 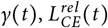 and *STIM*(*t*):

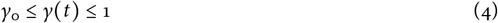

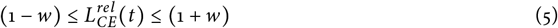

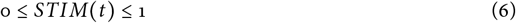

We implemented our problem as a Multiphase Optimal Control Problem to assure that CE shortening and lengthening occurred only once per full periodic cycle, which was done by inequality constraints on 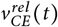:

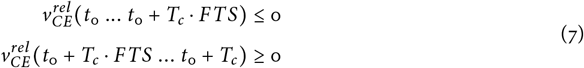

FTS was a numerical value (between 0.05 and 0.95) when FTS were imposed (see above). When FTS was not imposed, FTS was a control variable optimised for maximal AMPO.

When MTC length excursion was imposed, an extra inequality constraint was added:

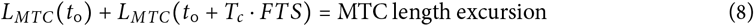

The dynamic, task and (in)equality constraints were implemented in CasADi (Andersson et al., 2019) and the cost function was minimised using a Direct Collocation method using a nonlinear interior point method (IPOPT, Wächter and Biegler, 2006). The optimal control solution was checked by running forward simulations of the model with a linear-interpolated 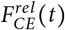 and *STIM* (*t*) as inputs to confirm that no relevant constraints were violated between the collocation points. This was done using SciPy’s ODE integrator with an Implicit Runge-Kutta method of Radau IIA family of order 5 (Hairer and Wanner, 1996) and an absolute and relative tolerance of 1e-6 and a maximum timestep of 1e-3. This yielded only very small differences between the states at the collocation points and those at the same time instance of the forward simulation for all optimisations.

### S4 Optimal CE length over time: influence of FTS and MTC length excursion

Using the Hill-type MTC model, we identified the optimal CE length over time for maximal AMPO without imposing any constraints on MTC length over time (in contrast to section Section 3.2.2, where MTC shortening and lengthening velocity were constant). When imposing cycle frequency, the resulting MTC length over time was reasonably similar to those when with constant MTC shortening and lengthening velocity (see Section 3.2.3 and Figure 10). However, when imposing FTS or MTC length excursion the shape of CE and MTC length over time dramatically changed.

For imposed FTS values below the optimal value, CE lengthening velocity was found to be almost zero at the onset of CE lengthening (Figure S2A). This infinitely low velocity at the start of CE lengthening was optimal for AMPO because it allowed the active state to decrease fully before lengthening occurred, thereby reducing the amount of negative mechanical work performed. On top of that, it allowed the active state to remain high throughout most part of CE shortening to enable substantial positive mechanical work. Towards the end of CE lengthening, CE velocity again approached zero, which made it possible to initiate muscle stimulation before CE shortening began. This resulted to a high active state at the onset of CE shortening to enhance positive mechanical work. For imposed FTS values above the optimal value, CE shortening velocity decreased over time at the end of CE shortening (Figure S2A). To avoid substantial negative mechanical work during the subsequent CE lengthening, muscle stimulation ceased well before the end of CE shortening. While CE could theoretically maintain a constant shortening velocity throughout the shortening phase, any additional positive mechanical work gained near the end was outweighed by the increased negative mechanical work. Thus, an almost isometric phase preceding CE lengthening is optimal for maximising AMPO. In sum, the CE length over time for imposed FTS values mostly results from the effect of the excitation dynamics.

For imposed MTC length excursions above the optimal value, CE shortening velocity was found to be higher at the beginning and end of the shortening phase compared to the middle part (Figure S2B). At the start and end of the shortening phase, only limited positive mechanical work could be produced due to the low active state and the suboptimal CE length (that is due to the CE force-length relationship). While CE could theoretically shorten at a constant velocity throughout the entire shortening phase, this would reduce the positive mechanical work during the middle part due to the effect of the force-velocity relationship. At the same time, little additional positive mechanical work could be achieved at the beginning and end of the shortening phase (due to the low active state and suboptimal CE length). Consequently, a non-constant CE shortening velocity — with CE shortening first decreasing and then increasing during the shortening phase — is optimal for maximising AMPO at imposed MTC length excursions above the optimal value. For imposed MTC length excursions below the optimal value, the opposite pattern was found: CE shortening velocity was found to be slower at the beginning and end of the shortening phase compared to the middle part (Figure S2B). As explained, only limited positive mechanical work could be produced at the beginning and end of the shortening phase due to the low active state. If CE would shorten with a constant velocity throughout the entire shortening phase, it would decrease the shortening distance during the middle part (at which the active state is high) and thereby reducing the positive mechanical work. As a result, a non-constant CE shortening velocity — with a slow shortening velocity at the end and beginning of the shortening phase — is optimal for maximising AMPO at imposed MTC length excursions above the optimal value. In sum, the optimal CE length over time for imposed MTC length excursions shows the intricate trade-off between the CE force-length relationship, CE force-velocity relationship and the and excitation dynamics.

**Figure S2:**
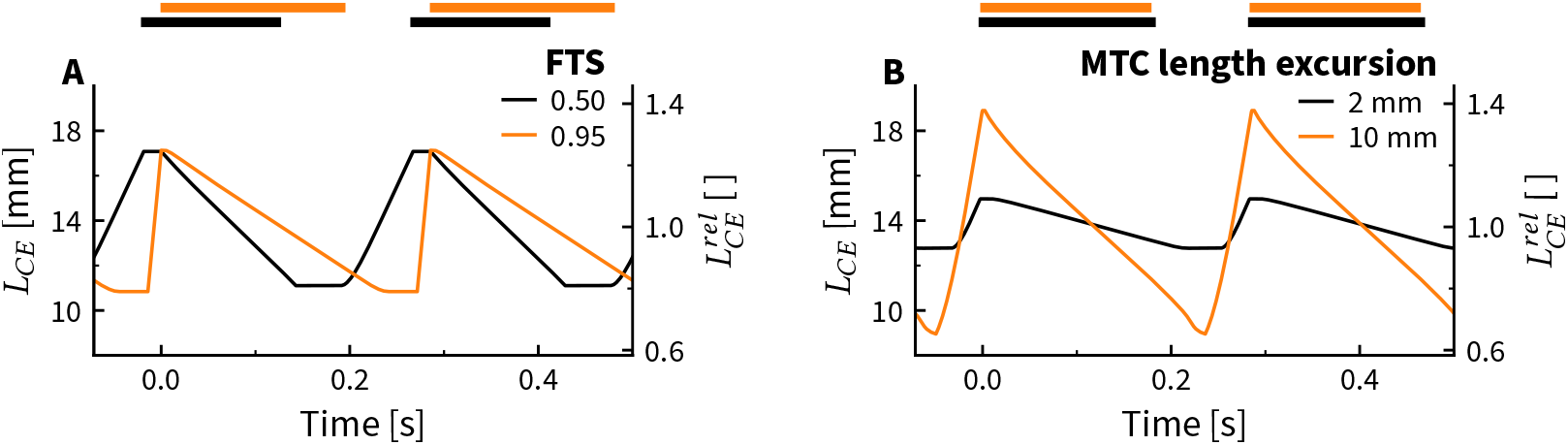
Predicted CE length over time for maximal attainable AMPO, shown for two imposed FTS values (A) and two imposed MTC length excursions (B), at a cycle frequency of 3.5 Hz for rat 1. In panel A, only cycle frequency and FTS were imposed, while in panel B only cycle frequency and MTC length excursions were imposed. Apart from these two constraints, CE and MTC length over time were completely unconstrained and thus the shape of CE length over time could be different between every combination of imposed cycle frequency and FTS (A) or imposed cycle frequency and MTC length excursion (B). CE stimulation was maximal during the period indicated by the coloured bars and off elsewhere.

**Figure S3:**
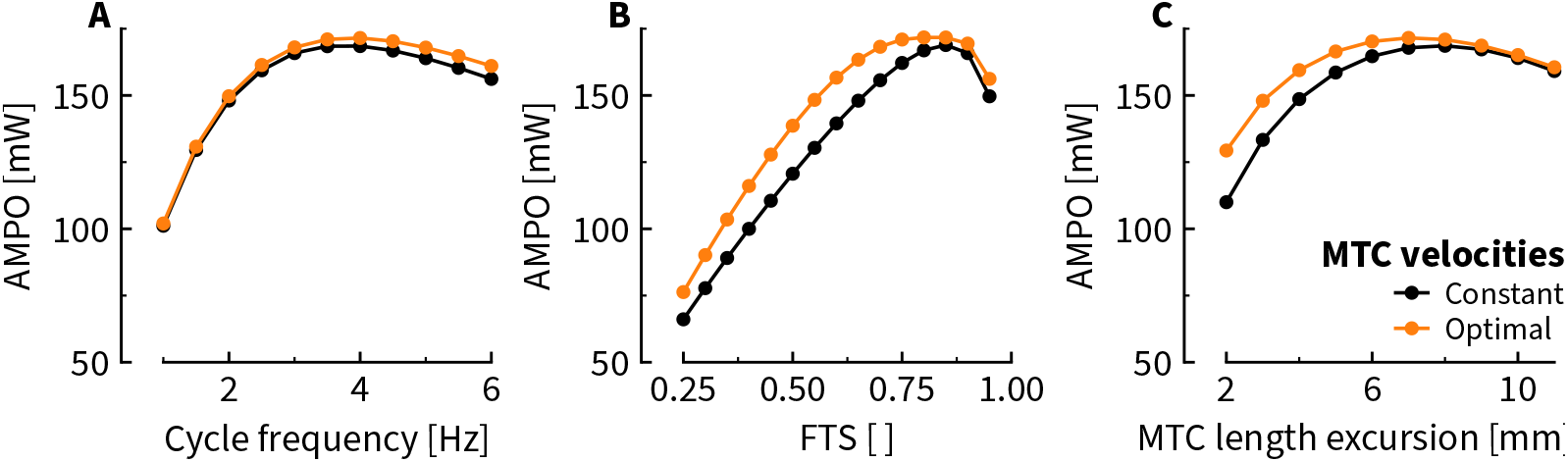
Predicted influence of cycle frequency (A), FTS (B) and MTC length excursion (C) on the maximum attainable AMPO. The maximal attainable AMPO — averaged across the three rats — slightly increased when there was no constraint on MTC length over time (orange lines), compared to SSCs with a constant MTC shortening and lengthening velocity (black lines).

